# eDNA metabarcoding provides scalable and continuous biodiversity monitoring across the tree of life

**DOI:** 10.64898/2026.02.12.705487

**Authors:** Till-Hendrik Macher, Robin Schütz, Jens Arle, Arne J. Beermann, Peter Haase, Jan Koschorreck, Henrik Krehenwinkel, Demetrio Mora, James S. Sinclair, Jonas Zimmermann, Florian Leese

## Abstract

Environmental DNA (eDNA) metabarcoding has the potential to substantially expand our knowledge of global biodiversity beyond that provided by conventional approaches. However, the degree to which eDNA data provides real ecological insight, rather than primarily reflecting environmental factors that affect eDNA shedding and degradation, remains unclear. Additional uncertainties arise in terms of cost-effectiveness and whether the price is worth any extra biodiversity information that is gleaned. Here, we established a high-resolution, bi-weekly eDNA time-series in Germany across aquatic and riparian habitats to quantify seasonal biodiversity dynamics, to relate eDNA to different potential environmental drivers, and to parametrize cost estimates. Over one year, eDNA metabarcoding detected more than 1,000 species across multiple trophic levels and primarily revealed real, taxon-specific seasonal patterns, in addition to some relationships to water temperature, discharge, and conductivity. Compared to historical records dating back to 1891, year-round eDNA monitoring increased reported species numbers by 2.4-fold for invertebrates, 2.2-fold for mammals, 1.7-fold for diatoms, 1.2-fold for fish and lamprey, and 1.03-fold for birds. Cumulative biodiversity estimates increased strongly with sampling frequency, demonstrating the value of eDNA for high-frequency time-series monitoring. Moreover, eDNA monitoring was highly cost-effective, providing more than twice the biodiversity information of many conventional surveys for one-sixth the cost, enabling scalable, high-resolution freshwater biodiversity assessments.

## Introduction

Biodiversity loss is one of the most pressing issues of the 21^st^ century ^1,2^. Addressing this loss requires comprehensive monitoring data to understand how biodiversity is changing and why. However, such biodiversity data are commonly lacking ^3^. For example, many biodiversity related legislations include monitoring components, but these suffer from incompleteness or insufficient sampling frequency to accurately characterize biodiversity dynamics. Current monitoring efforts also primarily use observational data and morphological specimen identification, introducing further limitations to the spatial and temporal extent of monitoring due to constraints of time and cost. This problem is further exacerbated in highly diverse and difficult to identify taxa, such as certain invertebrate groups. Additional biodiversity monitoring solutions are therefore essential for expanding our limited understanding of biodiversity change, ideally combining low-cost sampling with broad-scale and high-frequency monitoring, resulting in as close to real-time as possible ^4,5^.

Analysis of environmental DNA (eDNA), specifically that obtained through metabarcoding of freshwater samples, presents one possible solution for improved biodiversity monitoring ^6–8^. Owing to the relatively low sampling effort and recent advances in laboratory processing ^9^, eDNA metabarcoding has emerged as a potentially fast and low-cost monitoring method ^7,10,11^. Additionally, a single eDNA sample has the potential to yield biodiversity data on organisms across the tree of life, from bacteria to mammals ^12–15^. This data is not limited to freshwater taxa and can include non-aquatic species living in water bodies or interacting with the riparian zone such as mammals, amphibians, and birds ^16,17^. Such detections are possible because freshwaters gather and integrate inputs from surrounding terrestrial ecosystems. While freshwater eDNA has a proven potential for use in biodiversity monitoring ^18–21^, it remains uncertain how much new biodiversity data it provides compared to traditional methods, the degree of data reliability with respect to the inherent dynamics of rivers, and whether it can be feasibly integrated into broad-scale and high-frequency monitoring programs at an affordable cost ^6,22,23^.

To help determine this, we collected eDNA samples from a site in a restored river every two weeks for one year and conducted multi-taxa metabarcoding to identify as many species as possible across the tree of life. We quantified the amount of new biodiversity data provided based on the degree to which the eDNA data matched taxa recorded at and around the site for six major organism groups (fish, birds, mammals, freshwater invertebrates, terrestrial invertebrates, and diatoms). We assessed whether changes in eDNA species richness and read numbers for these taxonomic groups throughout the year reflected expected seasonal species changes (*biological explanation*) or were linked to changes in chemo-physical parameters (*chemo-physical explanation*). To test the biological explanation hypothesis, we determined whether taxon-specific changes in species composition and relative read abundance reflected known, phenological species shifts (e.g., seasonal temperature, fish spawning biology, migration patterns in birds). To test the alternative chemo-physical hypothesis, we analysed species relationships with discharge, conductivity and pH as key factors affecting the ‘ecology of eDNA’ ^24^, specifically eDNA dilution, transport and degradation ^25^. Lastly, to evaluate cost-effectiveness, we compared the costs of generating our results with those of traditional monitoring approaches in comparable settings and projected eDNA metabarcoding costs to an implementation scenario in which eDNA-based assessments complement monitoring under the EU Water Framework Directive (WFD). Our results demonstrate that eDNA metabarcoding is a non-invasive and efficient approach for tracking biodiversity across the tree of life, with strong potential to enhance both comprehensive biodiversity research and the cost-effectiveness of biological status assessments.

## Results

### Monitoring biodiversity across the tree of life

We conducted eDNA metabarcoding on 52 one-litre water samples collected from November 2020 to October 2021 (n=26) at the restored mouth of the River Lippe, Germany. In total, we detected 1,072 species, including 107 vertebrates, 775 invertebrates, and 190 diatoms. Among these, 41 were birds, 26 mammals, 40 fish, 425 freshwater invertebrates, 350 terrestrial invertebrates, and 190 diatom species (Fig. 1).

**Figure 1:**
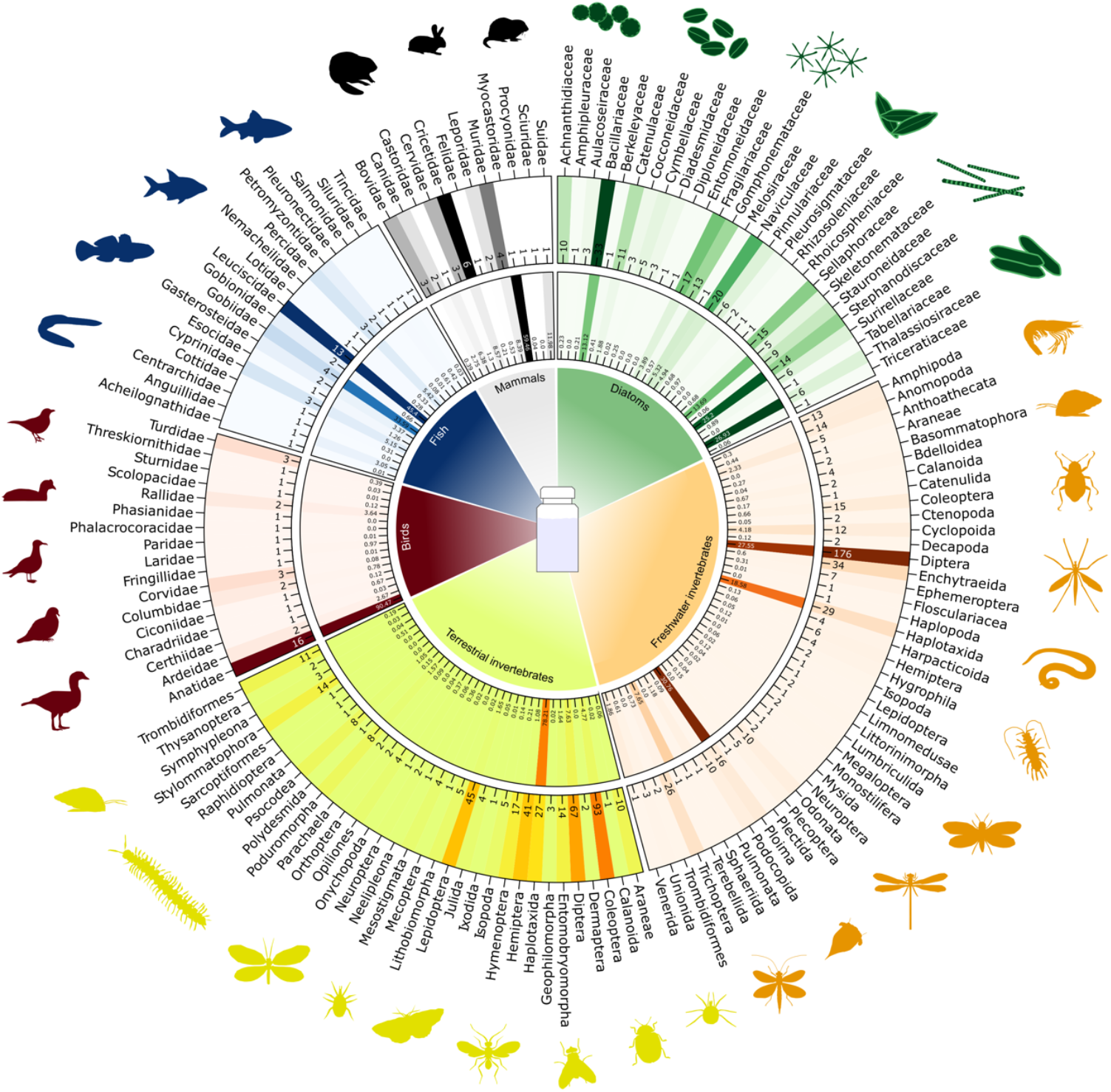
Biodiversity monitoring across the Tree of Life. The circular plot displays heatmaps of read proportions (inner circle) and the number of species (middle circle) detected for each taxon (outer circle), including diatoms, freshwater and terrestrial invertebrates, birds, fish, and mammals, derived from eDNA metabarcoding from water samples over the course of one year at the river Lippe mouth.

Among vertebrates, we detected 40 fish and lamprey species, with the cyprinid family Leuciscidae accounting for one-third of species (13) and nearly half of all fish reads (∼45%) (Table S1). Notably, we detected four highly invasive goby species (Gobiidae) that accounted for ∼34% of the total fish reads, with the round goby (*Neogobius melanostomus*) standing out as the dominant invasive alien species. Other notable families included carps and minnows (Cyprinidae) and perches (Percidae), alongside diadromous species such as the sea lamprey (*Petromyzon marinus*) and European flounder (*Platichthys flesus*). Beyond fish and lampreys, we detected 41 bird species, dominated by waterfowl of the family Anatidae (16 species of ducks, geese, and swans), which represented ∼40% of species and ∼90% of reads. Additional water birds included gulls (Laridae), rails (Rallidae), and cormorants (Phalacrocoracidae), while terrestrial bird detections included thrushes (Turdidae), crows (Corvidae), and pigeons (Columbidae). Among mammals (26 species), rodents (Rodentia) dominated. The family Cricetidae (voles and muskrats) alone represented nearly a quarter of species. Other terrestrial families included deer (Cervidae), hares and rabbits (Leporidae), true mice and rats (Muridae), and raccoons (Procyonidae). Semi-aquatic mammals were represented by the Eurasian beaver (*Castor fiber*) and the invasive nutria (*Myocastor coypus*), with the latter contributing ∼59% of mammal reads despite representing only ∼4% of species.

Detected invertebrates were primarily freshwater taxa (425 out of 775 species; Table S1). Insects (Insecta) dominated freshwater taxa (250 species, ∼57%), especially flies (Diptera; 176 species), including non-biting midges (Chironomidae), black flies (Simuliidae), and crane flies (Tipulidae). Classical bioindicator taxa were well represented, with 38 species of mayflies (Ephemeroptera), stoneflies (Plecoptera), and caddisflies (Trichoptera). Other aquatic groups included worms (Annelida, Nematoda, Nemertea, Platyhelminthes), freshwater polyps (Cnidaria), molluscs (Mollusca), and rotifers (Rotifera). Notably, rotifers of the order Ploima accounted for ∼31% of reads despite comprising only ∼4% of species. Terrestrial invertebrates comprised 350 species, again dominated by insects (244 species; ∼56%). The most diverse orders were beetles (Coleoptera; 93 species), flies (Diptera; 67), butterflies and moths (Lepidoptera; 45), and true bugs (Hemiptera; 41). Non-insect groups included earthworms (Haplotaxida and Acanthodrilidae; 15 species), land snails and slugs (Pulmonata and Stylommatophora; 15 species), and tardigrades (Parachaela; 2 species). Remarkably, reads were strongly skewed towards earthworms (Haplotaxida), which alone accounted for ∼78% of terrestrial invertebrate reads.

Lastly, we detected 190 diatom species (Bacillariophyta). The most diverse families were Bacillariaceae (33 species), Naviculaceae (20), and Fragilariaceae (17). However, read abundance was heavily dominated by only few families, with Stephanodiscaceae (14 species) and Thalassiosiraceae (6 species) together accounting for more than half of all diatom reads (Table S1).

### Comprehensiveness of eDNA biodiversity data

We evaluated data comprehensiveness by comparing the overlap between the taxa listed above to available historical records from our study site, dating back until 1891, for all taxonomic groups from the Global Biodiversity Information Facility (GBIF) ^26^, and from traditional monitoring of fish (2016 at 4 sites, 2019:4), freshwater invertebrates (2026:4, 2019:4, 2022:4), and diatoms (2018:2, 2020:3, 2021/2, 2022/1) conducted during the site restoration between 2016 and 2022. While the long-term historical baseline illustrates the full species pool ever recorded in the area, the more recent restoration monitoring data provide a realistic benchmark of the contemporary community against which eDNA performance can be assessed. Overall, eDNA-based monitoring detected slightly more total species over the course of one year than were historically reported in GBIF since 1891 (1,072 versus 1,029 species; Figure 2B). Specifically, eDNA metabarcoding recovered a substantial fraction of historically reported taxa, detecting 88.5% of fish species (23 out of 26), 45.5% of mammals (10 out of 22), 19.1% of birds (35 out of 183), 12.8% of all invertebrates (92 out of 733), and 14.8% of diatoms (12 out of 81) (Figure 2A). Beyond this overlap, eDNA metabarcoding significantly increased the number of reported species for diatoms (+68.73%), invertebrates (+48.79%), mammals (+42.11%), and fish (+39.53%), whereas the increase in birds was marginal (+3.17%). In comparison with the restoration monitoring datasets, eDNA metabarcoding detected 97.2% of fish species, 52.5% of aquatic invertebrates, and 20.4% of reported diatoms (Figure 2A). Here, the strongest relative increase in species detection was observed for aquatic invertebrates (+86.56%), followed by diatoms (+54.18%) and fish species (+12.20%).

**Figure 2:**
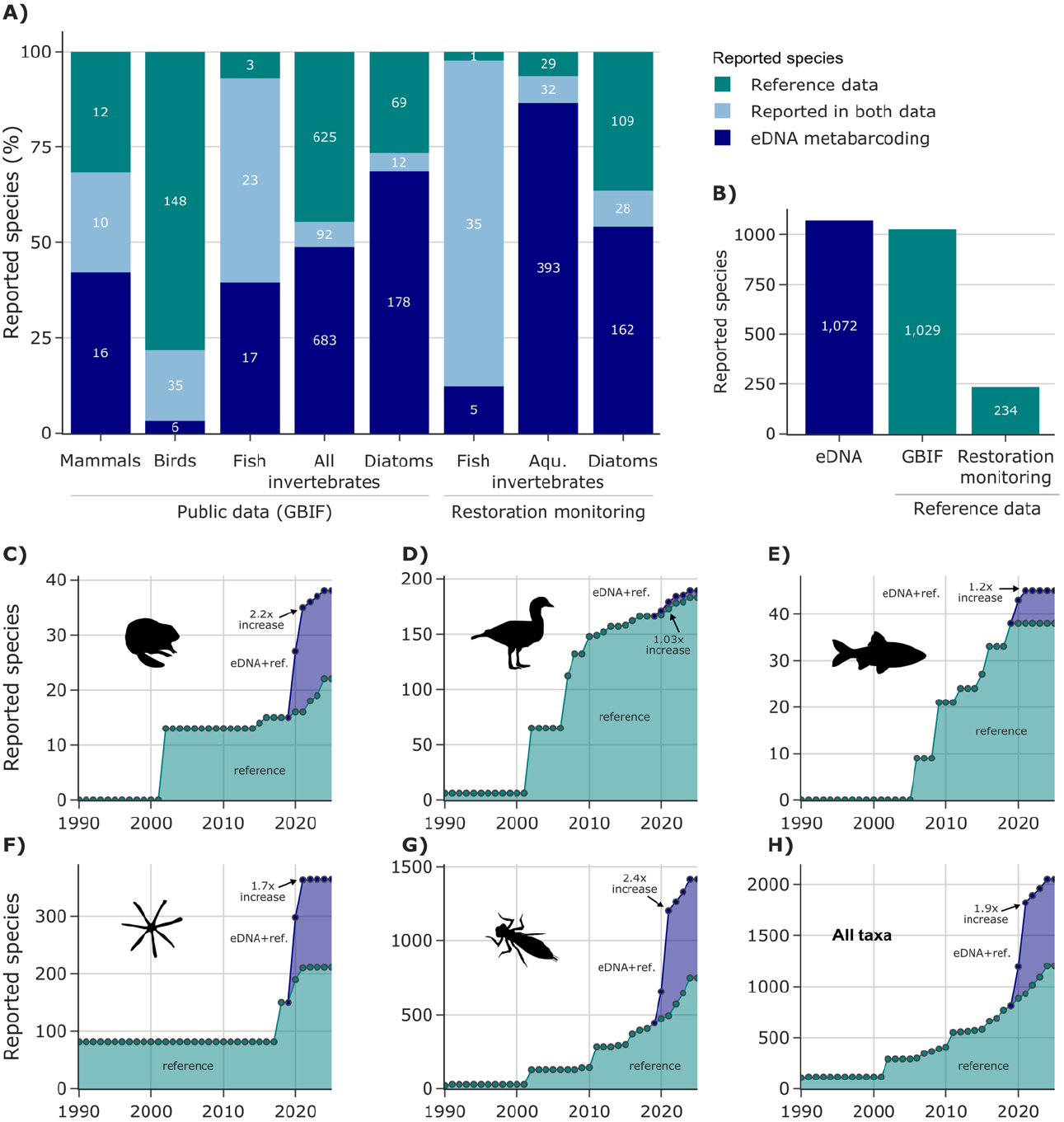
eDNA records in comparison to historical reports. (A) Comparison of species records from GBIF (mammals, birds, fish, invertebrates, and diatoms) and restoration monitoring (fish, freshwater invertebrates, and diatoms), combined referred to as reference data, with eDNA metabarcoding records. (B) Total number of species recorded across eDNA metabarcoding, GBIF, and restoration monitoring datasets. (C-H) Number of recorded species per organism group since 1990 based on eDNA metabarcoding and both reference datasets. The annual total number of recorded species is displayed once considering only the reference datasets (green) and then both the eDNA metabarcoding and reference datasets combined (blue). The factor of increase in reported species when including the eDNA dataset in the timeline is highlighted.

To further assess the contribution of eDNA to the monitoring data of the restored Lippe River mouth, we compared species richness in 2021 with and without the eDNA dataset (Figure 2 C-G). Invertebrates exhibited the highest increase in reported species (2.4x), followed by mammals (2.2x), diatoms (1.7x), and fish (1.2x). The number of reported bird species showed only a marginal increase (1.03x). Overall, the inclusion of the eDNA metabarcoding dataset increased the total number of reported species in 2021 by a factor of 1.9 (Figure 2H).

### Biological vs. abiotic drivers of eDNA dynamics

To assess whether observed seasonal dynamics is a result of the biology of species (*biological hypothesis*) or influenced by other factors that mobilise, transport or degrade eDNA (*chemo-physical hypothesis*) we quantified relationships between abiotic variables and both species richness and read abundances across all investigated taxa using Spearman rank correlations and generalized additive models (GAMs). To retain biologically meaningful relationships, analyses were restricted to statistically significant results (Spearman: p ≤ 0.05 and |ρ| ≥ 0.4; GAM: p ≤ 0.05 and pseudo-R^2^ ≥ 0.4).

Across taxa, water temperature (5.0-21.4 °C) showed the highest number of significant associations with biodiversity metrics, followed by conductivity (635-936 µS/cm) and water level (240-620 cm), which were also the only variables exhibiting significant temporal autocorrelation (Fig. S7). In contrast, pH (6.8-8.2) and precipitation (0-26.3 L m^-2^) showed comparatively few significant associations. These abiotic patterns form the basis for the taxon-specific seasonal analyses presented below.

Year-round eDNA monitoring revealed pronounced seasonal signals across most taxonomic groups, although their strength differed markedly (Fig. 3). Vertebrate communities showed comparatively moderate seasonal variation, whereas invertebrates and diatoms exhibited strong seasonal structuring.

**Figure 3:**
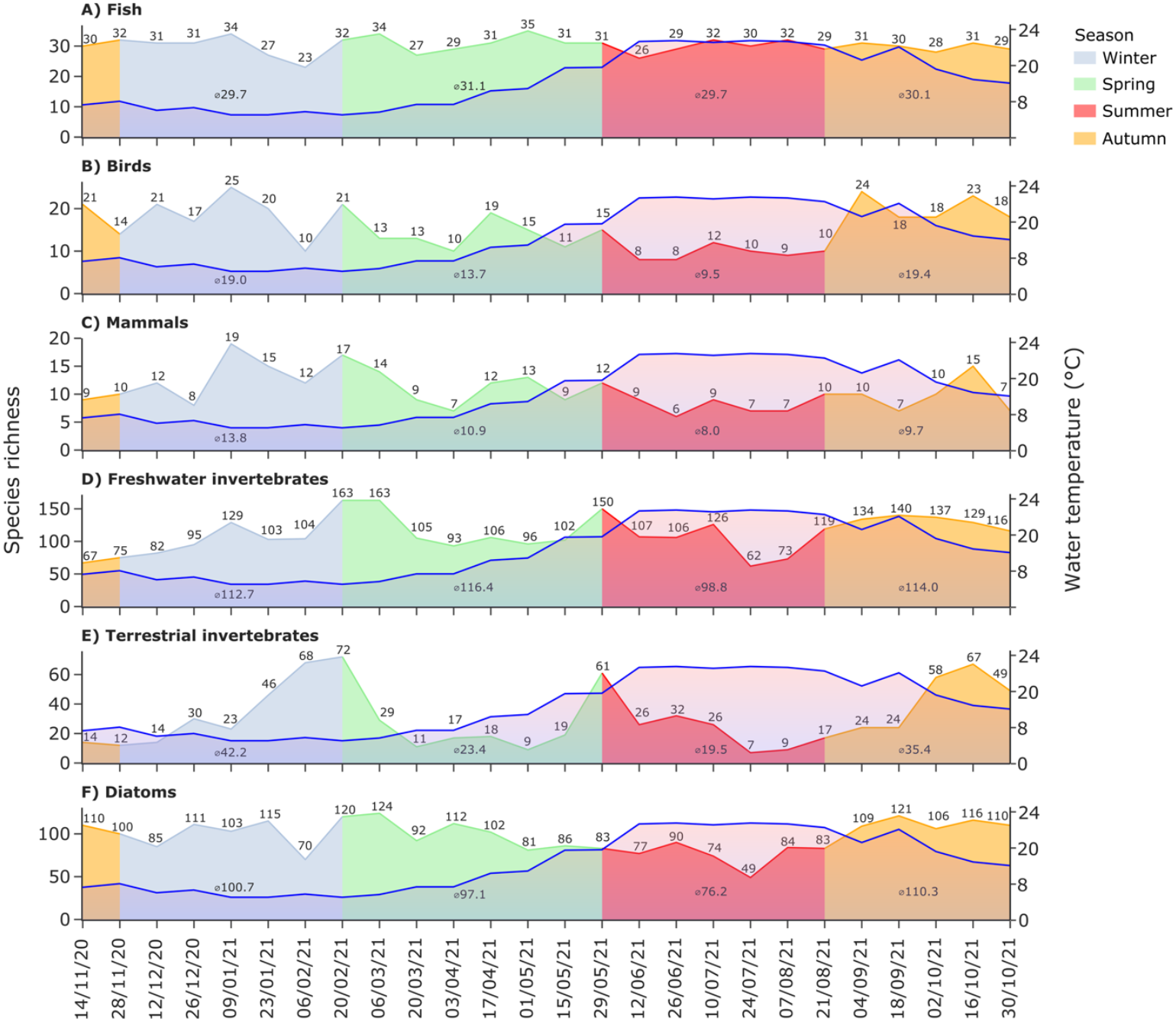
Seasonality of species richness. Number of observed species for fish and lamprey (A), birds (B), mammals (C), freshwater invertebrates (D), terrestrial invertebrates (E), and diatoms (F) over the course of one year. Exact species numbers are displayed for each sampling event. Additionally, the average number of species per season was calculated.

Specifically, fish species richness varied only weakly across seasons (23-35 species; Fig. 3A), and community composition differed modestly but significantly (ANOSIM: R = 0.11, p = 0.018; Fig. S6A). In line with the biological explanation hypothesis of seasonal patterns, observed changes in richness and relative read abundance were largely linked to spawning ecology rather than across-taxon patterns (overall richness, relative read abundance), with distinct temperature optima for cold, moderate, warm, and hot spawners (Fig. S2A). For example, eDNA read proportions of burbot (*Lota lota*) and northern pike (*Esox lucius*) peaked during winter, coinciding with their spawning periods, while GBIF records were concentrated in late summer, resulting in significant negative correlations between eDNA signals and GBIF occurrences (Fig. 4A).

**Figure 4:**
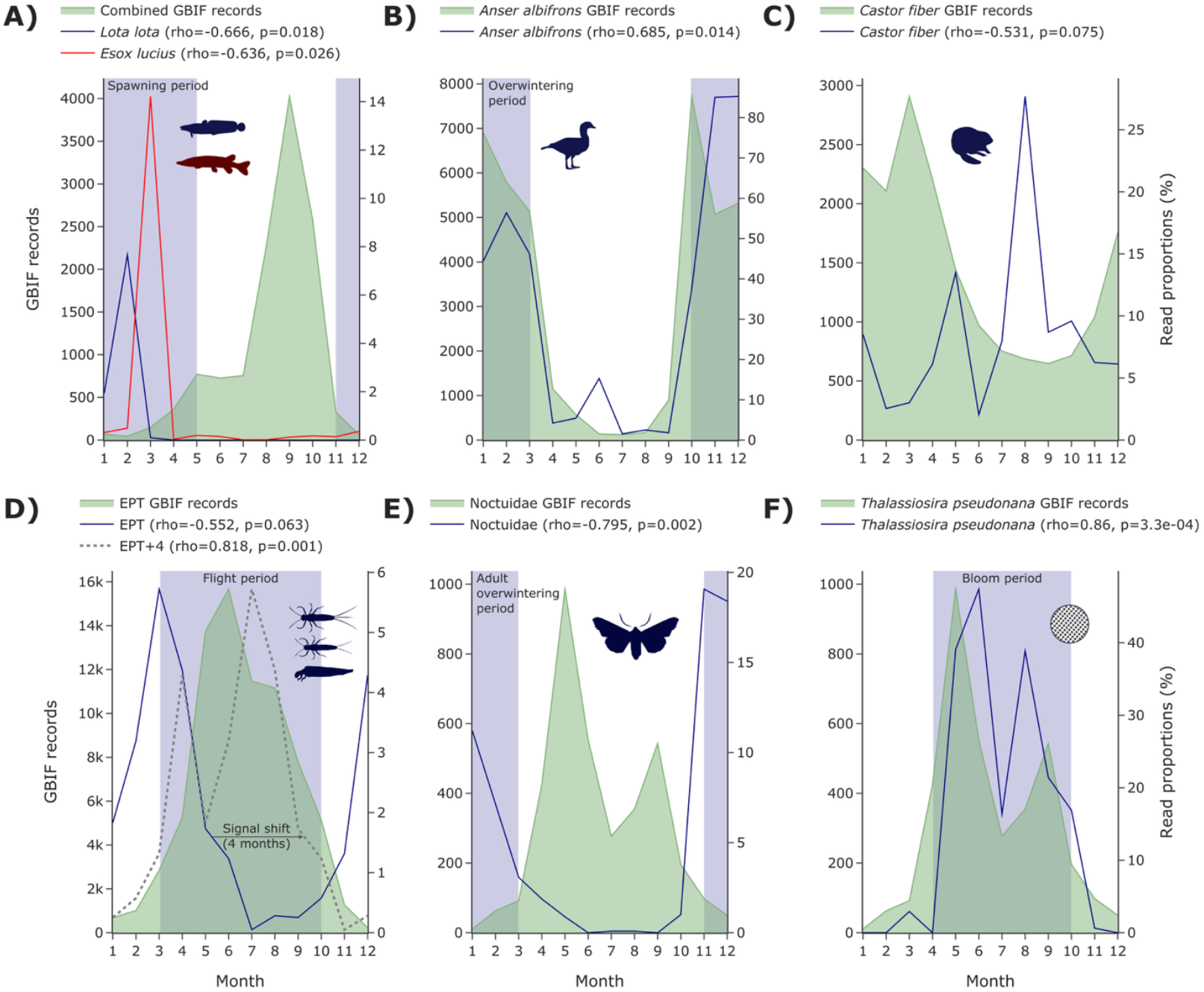
Examples of seasonal dynamics in eDNA signals compared to GBIF species occurrences. The number of GBIF records (filled green curve) and relative read proportions (blue, red, and grey dotted lines) per taxon are shown. (A) The winter-spawning fish species of burbot (*Lota lota*) and northern pike (*Esox lucius*) showed distinct eDNA peaked during their reproductive season in winter (blue background). (B) The white-fronted goose (*Anser albifrons*), a common winter visitor near the sampling site (blue background), exhibited overlapping temporal patterns between eDNA detections and GBIF observations. (C) The Eurasian beaver (*Castor fiber*) is present year-round, which is consistently reflected in the eDNA data. (D) EPT taxa (orders Ephemeroptera, Plecoptera, and Trichoptera), important ecological indicators, were primarily detected during winter months in eDNA samples. Their signal preceded GBIF records by approximately four months, which mostly represented occurrences during the terrestrial flight period (blue background). (E) Noctuid moths (Noctuidae), known to include overwintering species active on mild winter nights (blue background), showed higher detectability in eDNA during winter months, contrasting to increased GBIF reports during summer. (F) The common diatom *Thalassiosira pseudonana* showed matching eDNA signals and GBIF occurrences both peaking twice, in late spring and early autumn (blue background).

Bird communities displayed stronger seasonal variation, with species richness ranging from eight species in summer to 25 species in winter (Fig. 2B), and clear seasonal clustering in community composition (ANOSIM: R = 0.46, p = 0.001; Fig. S5B). Seasonal patterns reflected migratory behaviour, with migratory species showing increased detections at lower temperatures (Spearman ρ = 0.712), whereas stationary species exhibited higher read proportions at warmer temperatures (Spearman ρ = -0.712) (Fig. S2B). The white-fronted goose (*Anser albifrons*) exemplified this pattern, with winter peaks in eDNA detection strongly correlated with GBIF records (Fig. 4B).

Mammal communities showed moderate seasonal variation in species richness (7–19 species; Fig. 3C) but no significant seasonal shifts in community composition (ANOSIM: R = 0.09, p = 0.066). Most species were detected consistently throughout the year, such as the Eurasian beaver (*Castor fiber*) (Fig. 4C).

In contrast, freshwater invertebrates and diatoms exhibited pronounced seasonal dynamics. Freshwater invertebrate richness peaked in spring and declined in summer (Fig. 3D), with strong seasonal clustering in community composition (ANOSIM: R = 0.56, p = 0.001; Fig. S7A). EPT (Ephemeroptera, Plecoptera, Trichoptera) taxa were predominantly detected during winter, and although eDNA signals were not synchronous with GBIF records, they preceded reported occurrences by approximately four months and correlated strongly after temporal shifting (Fig. 4D). Terrestrial invertebrates showed similar but weaker seasonal structuring (ANOSIM: R = 0.36, p = 0.001; Fig. S7B), with noctuid moths displaying increased winter detectability in eDNA and an inverse relationship with GBIF occurrences (Fig. 4E).

Diatom communities also showed strong seasonal turnover, with highest richness in autumn and lowest richness in summer (Fig. 3F) and pronounced seasonal clustering (ANOSIM: R = 0.53, p = 0.001; Fig. S7C). While species richness was only weakly associated with abiotic variables, read proportions responded strongly to seasonal temperature variation. For example, the diatom species *Thalassiosira pseudonana* showed closely matching seasonal patterns in eDNA detections and GBIF records (Fig. 4F).

Abiotic influences on biodiversity metrics were strongly taxon- and trait-dependent (Fig. S8, S9). Across all tested taxon–abiotic variable combinations, the number of significant associations was comparatively low for species richness (Spearman’s ρ n = 759; GAM R^2^ n = 31), whereas read proportions exhibited substantially stronger and more frequent correlations (Spearman’s ρ = 1516; GAM R^2^ = 672). Among vertebrates, species richness was only weakly associated with abiotic factors (Spearman’s ρ = 141; GAM R^2^ = 8), while read proportions showed markedly higher sensitivity (Spearman’s ρ = 280; GAM R^2^ = 152). In contrast, invertebrate read proportions were strongly influenced by multiple abiotic drivers, particularly water temperature (Spearman’s ρ = 328; GAM R^2^ = 120), conductivity (Spearman’s ρ = 225; GAM R^2^ = 104), and water level (Spearman’s ρ = 226; GAM R^2^ = 69). Similarly, diatom read proportions exhibited strong responses to abiotic conditions, with water temperature emerging as the dominant driver (Spearman’s ρ = 172; GAM R^2^ = 58).

### eDNA cost estimates

To evaluate the cost effectiveness of multi-taxon eDNA metabarcoding, we quantified the expenses associated with processing water samples across different taxonomic groups. The total cost of metabarcoding 52 water samples targeting birds, fish, mammals, invertebrates, and diatoms -including laboratory consumables, reagents, sequencing, and personnel time (non-commercial setting) for two PCR replicates - amounted to €11,938.26, corresponding to €229 per sample (Table S17). The analysis of vertebrates (birds, fish, and mammals) costed €65 per sample, which also accounted for the invertebrates, summing up to €3,396.09 (Fig. 5A). Owing to the higher sequencing requirements associated with longer barcode fragments, diatom analyses incurred higher costs, totalling €5,146.09, or €99 per sample.

**Figure 5:**
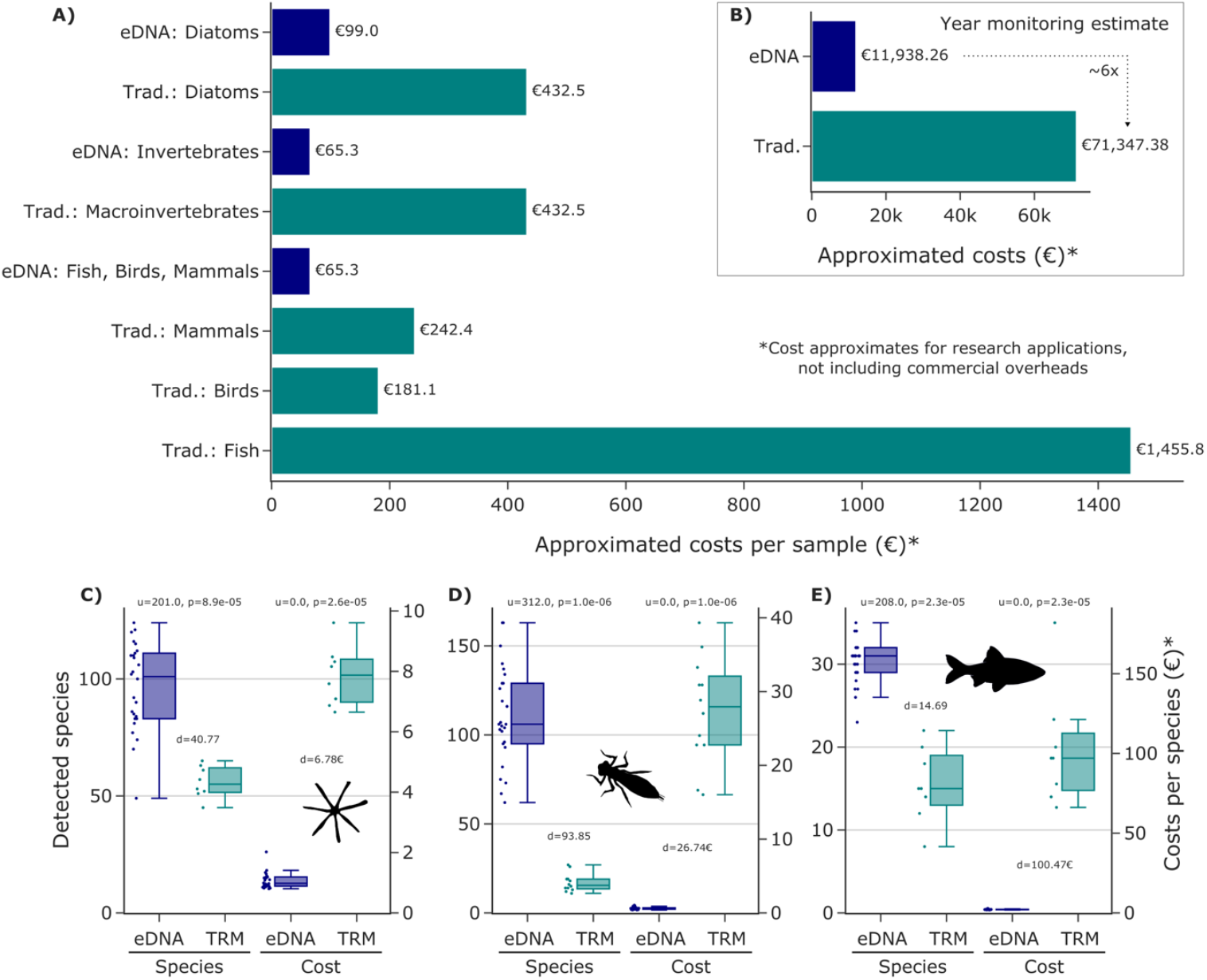
Cost comparison and species detection efficiency of eDNA metabarcoding versus traditional (morpho-taxonomic) monitoring. (A) Per-Sample costs: Estimated costs per sample for the eDNA metabarcoding approach and traditional (morpho-taxonomic) approach, both targeting vertebrates (birds, fish, and mammals), invertebrates, and diatoms. (B) Total Annual Costs: Approximate total costs of both monitoring approaches over an exemplary one-year period with biweekly sampling. For the eDNA approach, costs were calculated for 52 samples (including two field replicates), while traditional costs were based on 26 sampling events. Labour costs were standardized at 60.35 € per hour. (C-D) Species detection and cost efficiency: The number of species detected per sample was compared between eDNA-based monitoring from this study and the official restoration monitoring (TRM) conducted between 2016 and 2022, targeting diatoms, freshwater invertebrates and fish. Based on these species counts and the respective cost estimates, the cost per detected species (€) was calculated for each method. Statistical differences between methods were assessed using Mann-Whitney U tests.

**Figure 6:**
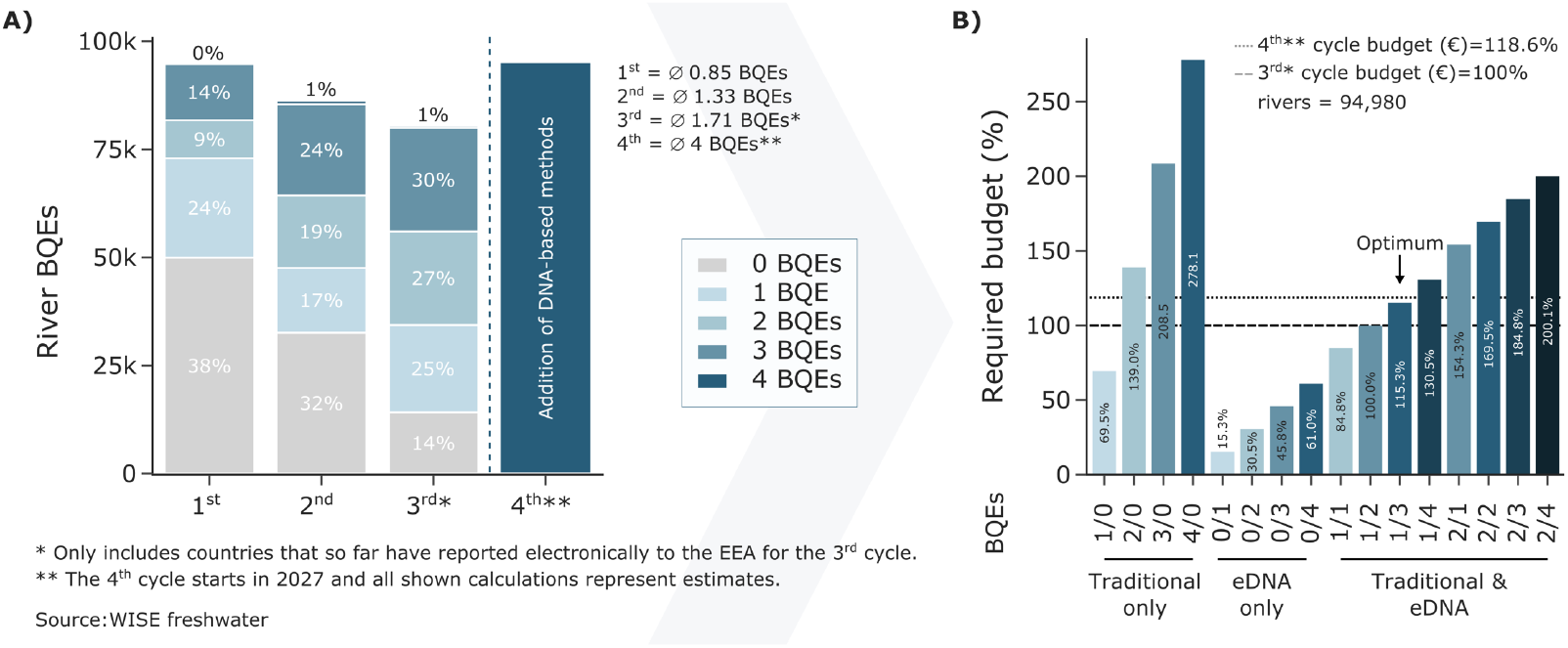
Potential analysis of (e)DNA metabarcoding in regulatory monitoring. Illustration of a cost estimation to assess the implementation potential of (e)DNA metabarcoding as a complementary biomonitoring method within the Water Framework Directive (WFD). (A) The number of river biological quality elements (BQEs) assessed per stream has been steadily increasing from 0.85 BQEs to 1.71. BQEs within the first three WFD cycles. DNA-based methods have the potential to help fill these gaps and improve WFD monitoring to up to four assessed BQEs within a similar budget. (B) The costs for river biomonitoring over the last three WFD cycles were estimated by multiplying the average cost per traditional sample ^27^ by the number of Biological Quality Elements (BQEs) assessed per river. The budget for the 4^th^ cycle was projected based on previous budgets. Cost estimates for 16 potential BQE assessment scenarios, including traditional-only (X/0), (e)DNA-only (0/Y), and combined approaches (X/Y), were calculated. The optimum was estimated with one traditionally assessed BQE and three BQEs assessed with eDNA metabarcoding, requiring 115.3% € of the 3^rd^ cycle budget.

In comparison, the total estimated costs for traditional monitoring conducted over one year, which does not require the extra replicate used for metabarcoding and so equates to 26 sampling events covering fish, birds, mammals, freshwater invertebrates, and diatoms, amounted to approximately €71,347.38 (Figure 5B, Table S17). Specifically, costs were estimated for electrofishing at €37,850.28, bird visual census at €4,707.30, mammal camera trapping at €6,302.40, freshwater invertebrate bulk sampling at €11,243.70, and diatom assessments at €11,243.70. We note that terrestrial invertebrates were not included in these cost estimates, as they are not routinely covered in traditional monitoring programs. Thus, eDNA metabarcoding as implemented here can be implemented for one-sixth the cost of traditional monitoring approaches (Figure 5B).

To directly compare effectiveness, we calculated costs per detected species at the sample level for both the eDNA-based assessment and the restoration monitoring data, using species richness per sample for fish, freshwater invertebrates, and diatoms (Fig. 5C-E). This resulted in paired, sample-by-sample estimates of cost per species for each method. Across all three organism groups, and under the chosen cost setting, eDNA metabarcoding yielded substantially lower cost-per-species values than traditional restoration monitoring for fish (U = 0, p = 2.3 x 10^-5^), freshwater invertebrates (U = 0, p = 1.0 × 10^-6^), and diatoms (U = 0, p = 2.6 ×10^-5^).

## Discussion

Here, we demonstrated biweekly sampling at a restored river mouth as a proof-of-concept to demonstrate the scalability and sensitivity of eDNA metabarcoding, detecting over 1,000 species across the tree of life from water samples. Our sampling design highlights the scalability of eDNA metabarcoding, enabling monitoring strategies that can flexibly balance temporal resolution and spatial coverage.

Beyond taxonomic coverage, eDNA captures biologically meaningful seasonal dynamics reflecting species-specific life-history strategies. Seasonal variation in richness and read abundance was highly taxon-specific, with pronounced signals in taxa with discrete reproductive or migratory periods. Such asynchronous trajectories are difficult to reconcile with purely abiotic explanations, which act broadly across taxa, and instead align with established ecological traits such as spawning and migration timing. While abiotic factors modulate eDNA detectability, they do not appear to be the primary driver of the observed seasonal patterns. Across the fish community, multiple species showed seasonally elevated eDNA signals consistent with known reproductive phenologies; as illustrative examples, winter-spawning fishes such as burbot (*Lota lota*) and northern pike (*Esox lucius*) exhibited elevated eDNA signals in cold months, consistent with increased DNA release during reproduction rather than biomass changes ^28,29^. This supports previous findings that eDNA integrates biological activity as well as presence, necessitating life-history-aware interpretation ^30^. Phenology-driven eDNA patterns may diverge from conventional monitoring records collected in other seasons, highlighting the importance of aligning sampling with biological rhythms ^31^.

Seasonal eDNA signals also correlated with historical occurrence data. For example, white-fronted geese (*Anser albifrons*) exhibited strong temporal concordance with historical overwintering records ^32^. Similar alignment was observed for key freshwater EPT taxa (mayflies, stoneflies, and caddisflies) which overwinter as larvae and emerge as adults in spring, showing peak eDNA signals in winter while GBIF adult records peak in summer ^18,33^. Diatoms such as *Thalassiosira pseudonana* also displayed bimodal bloom dynamics consistent with both eDNA and occurrence records. Terrestrial species that interact sporadically with water generally showed less consistent eDNA patterns. For example, an overwintering noctuid moth ^34^ displayed an inverse pattern compared to GBIF records: eDNA detectability was higher in winter, while traditional observations peaked in summer. This discrepancy likely reflects reduced overall invertebrate activity in winter - boosting the relative eDNA signal - and observer biases in GBIF, which favour reporting during warmer months. Collectively, these examples demonstrate that eDNA metabarcoding can reveal fine-scale, quantitative biodiversity patterns across ecological traits and wet-dry cycles ^35^, though interpreting quantitative eDNA signals may require approaches that differ from traditional monitoring data.

Higher temporal resolution is increasingly important under global environmental change, where subtle and taxon-specific responses may be missed by infrequent sampling, regardless of the identification method employed ^36,37^. In this study, intensive temporal sampling is used to demonstrate the capacity of eDNA metabarcoding to capture such dynamics, rather than to define a fixed monitoring frequency. Separately, eDNA metabarcoding offers broad taxonomic coverage at relatively low cost, enabling scalable, repeatable, tree-of-life assessments across animals, plants, fungi, and microbes ^38^. While false negatives are inherent to eDNA-based surveys ^39^, no monitoring approach is exhaustive; morpho-taxonomic surveys similarly face limitations, including inconsistent taxonomic resolution, observer bias, and overlooking of cryptic taxa ^40,41^. Complementary and integrative frameworks are therefore essential. Consistent with this, our results reveal limited overlap between eDNA and traditional approaches for invertebrates, diatoms, and birds, underscoring that each method captures distinct components of biodiversity. While eDNA data are currently retrofitted into conventional indices ^41^, the development of eDNA-specific assessment frameworks that integrate abiotic conditions, land use, and ecological traits would enable more objective and taxonomically inclusive evaluations, including microbial communities largely excluded from traditional monitoring ^6,42^. Our data provide further evidence that high-frequency, year-round monitoring at ecologically sensitive sites can reveal phenological patterns and detect invasive species, while simultaneously also offering potential for pathogen surveillance and early warnings of biodiversity loss ^43–45^. Species detectability varies seasonally, highlighting the risk of misestimating biodiversity when relying on single time-point surveys. In this context, automated “biodiversity weather stations” could become transformative early-warning tools ^46^.

While continuous monitoring may be appropriate at critical nodes such as river mouths, year-round sampling is not feasible across large spatial programmes such as those implemented under the WFD. We therefore advocate spatially and temporally integrated monitoring strategies, in which eDNA metabarcoding complements conventional approaches rather than replacing them. Our cost projections indicate that a hybrid strategy, e.g. combining one traditionally assessed BQE with three eDNA-based BQEs derived from a single water sample, would enable the simultaneous assessment of four BQEs per river within the projected budget of the next WFD cycle, and with only little additional costs eDNA assessments can be added in addition. Although ecological status classes (ESCs) were not assigned in this study and an intercalibration is of eDNA-based assessment is still pending ^41^, the targeted sampling of WFD-relevant BQEs demonstrates the technical and economic feasibility of integrating eDNA metabarcoding into regulatory monitoring workflows. An initial phase of parallel traditional and eDNA-based BQE assessments is recommended to enable robust intercalibration across BQEs and validation of eDNA-based quantification for legislative monitoring. Although eDNA metabarcoding does not recover identical taxonomic assemblages compared to traditional methods and quantitative interpretation remains under development, its non-invasive nature substantially reduces harm to assessed species while offering a cost-effective and integrative approach to biodiversity assessment. Moreover, a single eDNA or BQE sampling effort can provide information on additional taxa beyond the targeted BQEs: for example, fish eDNA metabarcoding also detects birds and mammals, and freshwater invertebrate assessments capture terrestrial invertebrates. While these additional data do not yet fulfil formal criteria for inclusion in legal monitoring programs, they are obtained without any additional effort, increasing the overall value of each survey. This integrative potential extends the relevance of eDNA monitoring beyond the WFD to international frameworks such as the Kunming-Montreal Global Biodiversity Framework and the United Nations Sustainable Development Goals ^6^.

We therefore propose a three-tiered monitoring strategy: (i) broad, holistic eDNA surveys from single water samples to identify large-scale spatial and temporal trends; (ii) targeted morpho-taxonomic assessments to quantify biomass and abundance and maintain continuity with long-term datasets, benchmark, validate and intercalibrate eDNA data sets; and (iii) high-frequency time-series eDNA monitoring at critical nodes to resolve seasonal dynamics and provide early-warning signals. Traditional methods remain indispensable, but eDNA substantially enhances monitoring capacity by expanding taxonomic coverage, improving temporal resolution and increasing cost efficiency. Integrating these complementary tiers will support resilient and responsive monitoring systems capable of addressing current and future biodiversity challenges.

### Outlook

In summary, eDNA metabarcoding offers a scalable addition to existing biodiversity monitoring frameworks, enabling earlier and broader detection of ecological change. The next challenge lies in integrating these approaches into regulatory monitoring schemes, standardizing analytical workflows, and developing quantitative frameworks that translate molecular signals into management-relevant indicators. Achieving this integration will allow monitoring programs to build upon established assessment frameworks by incorporating broader taxonomic coverage and higher temporal resolution, ultimately strengthening biodiversity monitoring and management.

## Methods

Methodological details are provided in Supplementary Material 1.

### eDNA metabarcoding

Water samples were collected biweekly from November 14, 2020 to October 30, 2021 (26 sampling events) at the restored mouth of the River Lippe (Wesel, Germany), a site previously subjected to extensive anthropogenic modification. On each sampling date, two water samples were taken 10 minutes apart around noon and filtered on site using sterile 0.45 µm PES filters (Whatman, England). Three field blanks were included as negative controls. DNA extraction was performed directly from the filters using Proteinase K and the NucleoMag Tissue Kit (Macherey Nagel), with four extraction blanks included. We amplified eDNA using three primer sets: tele02 for vertebrates (12S, ∼167 bp) ^47^, fwhF2/fwhR2n for invertebrates (COI, 205 bp) ^48^, and Diat_rbcL_708F/R3 for diatoms (rbcL, 312 bp) ^49^. A two-step PCR protocol with two technical replicates per sample was used, followed by size selection and normalization with magnetic beads ^9,50^. Dual-indexed libraries were generated and pooled primer-wise. Libraries for tele02 and fwhF2/fwhR2n were sequenced on the Illumina HiSeq (2×150 bp, Macrogen, Europe), while the rbcL library was sequenced on a MiSeq (2×250 bp, CeGat, Germany).

### Bioinformatics

Demultiplexed FASTQ files were processed using the APSCALE-GUI v1.1.6 pipeline ^51^, with each primer dataset handled separately under default settings (see Supplementary Material 1). For tele02 and fwh2 data, OTUs were clustered at 97% similarity, a threshold appropriate for species-level resolution in fish and macroinvertebrates ^52,53^. The *rbc*L dataset was denoised into ESVs (alpha = 2), commonly used in diatom assessments ^49,54^.

Taxonomic assignment was conducted with primer-specific approaches: tele02 and rbcL sequences were processed using APSCALE-blast (https://github.com/TillMacher/APSCALE_blast) against Midori2 ^55^ and diat.barcode v11.1 ^56^, respectively. fwh2 OTUs were assigned via the BOLDigger pipeline ^57^. Assignments for tele02 and *rbc*L were filtered using APSCALE-blast (see Supplementary Material 1) and verified by taxonomic experts.

OTU and ESV tables were converted into TaXon tables (Tables S2–S4) using TaxonTableTools v1.4.7 ^58^. Technical PCR replicates were merged, retaining only taxa found in both replicates. Reads from negative controls were subtracted to reduce potential contamination. The datasets were filtered by taxonomic groups: tele02 (fish, lampreys, other vertebrates), fwh2 (macroinvertebrate phyla), and rbcL (Bacillariophyta). fwh2 taxa were further classified into freshwater and terrestrial groups by experts from LANUK. OTUs/ESVs with ≥97% similarity to a reference sequence but lacking species-level resolution were manually validated using GBIF occurrence data (GBIF.org, 2024). Field replicates were merged per date, and all downstream analyses were based on the resulting filtered tables (Tables S5– S10).

### Statistical analysis

To provide an overview of alpha biodiversity, species richness and eDNA metabarcoding read proportions for diatoms, freshwater invertebrates, birds, fish, and mammals were calculated and visualized using a circular plot. Seasonal patterns were examined by plotting richness over time and calculating seasonal averages per taxonomic group.

Correlations between biodiversity metrics and abiotic factors were then assessed. Fish and lampreys were categorized by spawning temperature preference using StoreFish 2.0 ^59^, while birds were grouped as migratory or stationary ^60,61^. Mammals were classified by habitat and hibernation strategy. Trait data for freshwater invertebrates (feeding type, life mode, locomotion) were obtained from freshwaterecology.info v8.0 ^62^, and diatoms were grouped by life mode. Terrestrial invertebrates were excluded from trait-based analyses.

Abiotic factors included water temperature, pH, conductivity (all measured in situ), water level (provided by EGLV), and precipitation (72-hour average prior to sampling; meteostat.net, accessed 22 February 2024). First, spearman’s rank correlation implemented in the spearmanr() function scipy ^63^ was used to examine associations between biodiversity metrics (richness and read proportions) and abiotic variables across taxonomic ranks and trait-based groups. Only significant results (p ≤ 0.05) with sufficient sample presence (≥ 4) and richness (≥ 4 species) were retained (Tables S12-S13). Similarly, generalized additive models (GAM) with a Poisson distribution and a “log” link function implemented by the gam() function in the R package mgcv ^64^ were used. Where overdispersion was detected, we corrected the standard errors using a quasi-GAM model. Only significant results (p ≤ 0.05) with pseudo-R^2^ values ≥ 0.4 and with sufficient sample presence (≥ 4) and richness (≥ 4 species) were retained (Tables S14– S15).

Community composition was assessed using principal coordinate analysis (PCoA; skbio python library, ver. 0.7.0) based on Jaccard distances for each taxonomic group, with samples grouped by season. ANOSIM tested for seasonal differences. Beta diversity (turnover and nestedness) was quantified pairwise and visualized using density plots, with group-wise averages also reported (Figures S4–S5).

Furthermore, one notable taxon for each of the six investigated taxonomic groups was assessed individually. For each taxon, the average relative read proportions per month were calculated. Then, occurrence data were downloaded from GBIF and the number of occurrences per month recorded in Germany for *Lota lota* and *Esox lucius* ^65^, *Anser albifrons* ^32^, *Castor fiber* ^66^, EPT taxa ^67^, and Noctuidae ^68^ as well as Europe-wide for *Thalassiosira pseudonana* ^69^ were calculated. Correlations were tested using Spearman’s rank correlation between the eDNA read proportions and GBIF occurrences.

### Method comparison

To compare eDNA metabarcoding with existing biodiversity data, species records from the GBIF database ^26^ and official restoration monitoring (EGLV, 2016–2022) were compiled for fish and lamprey, mammals, birds, freshwater invertebrates, and diatoms (Table S16). GBIF records ranged from 1891 to 2025. Species overlap among datasets was assessed to identify unique and shared taxa. Additionally, temporal trends in species richness were analysed by plotting cumulative species numbers since 1990 under two scenarios: (1) reference data only (GBIF + restoration monitoring) and (2) including reference combined with the eDNA dataset. Increases in reported species after 2020 were compared between timelines.

### Cost comparison

Per-sample costs for the eDNA metabarcoding workflow (this study) were calculated and compared with estimated costs for traditional morpho-taxonomic monitoring. Both material and labour costs (at 60.35 €/h) ^70^ were considered under academic/research settings, excluding capital expenditures. Estimates were based on 52 eDNA samples and 26 traditional sampling events. Costs were broken down by organism group and monitoring phase (sampling, lab work, analysis) and summarized in Table S17. Traditional monitoring costs were estimated per sample following established protocols. Bird monitoring assumed a single observer conducting a 3 h visual census. Fish monitoring followed electrofishing protocols including material upkeep ^71^ and three scientists sampling for 8 h. Mammal monitoring was based on camera trap upkeep and one scientist spending 2 h in the field and 2 h screening images. Macroinvertebrate and diatom monitoring included costs for sampling and preparation, WFD-compliant field sampling by two scientists (2 h each), and 3 h of morphological identification by one scientist. These calculations should be regarded as estimates and will differ between protocols. Lastly, the number of species detected per sample and the cost per detected species were calculated for diatoms, freshwater invertebrates, and fish. Method differences were statistically assessed using the Mann-Whitney U test.

### Application potential in regulatory monitoring

To evaluate the scalability of eDNA metabarcoding in EU Water Framework Directive (WFD) monitoring, we compared costs across 16 assessment scenarios (traditional-only, eDNA-only, and hybrid) for 94,980 rivers. Cost estimates were based on EU member state reports (Belgium, Netherlands, Slovakia, Spain) ^27^ and our own calculations (Table S18). The total budget for BQE assessments in the WFD’s three completed cycles was estimated using the number of rivers assessed with 1-4 Biological Quality Elements (BQEs) from the EEA’s WISE database. Projected costs for the 4th cycle (118.6% of 3rd cycle) were used to benchmark eDNA-based scenarios, which were visualized as relative proportions of the 3rd cycle budget. Finally, an implementation roadmap was developed, outlining three phases for integrating (e)DNA methods into regulatory frameworks: (1) method development, (2) validation and intercalibration, and (3) regulatory adoption.

## Acknowledgements

This study was conducted as part of the GeDNA project, funded by the German Federal Environmental Agency (FKZ 3719242040). We also thank the Emschergenossenschaft and Lippeverband (EGLV), with special appreciation to Gunnar Jacobs, for facilitating the collection of eDNA samples and for their support and feedback during the data analysis. We extend our gratitude to Susanne Schüler (LANUV) for her meticulous curation of the macroinvertebrate taxa list. TM was funded by the dbDNA project (German Federal Environment Agency: FKZ 3722232010). RS was funded by TrendDNA project (German Federal Environment Agency: FKZ 153606).

